# Eccentric cycling enhances primary motor cortex excitability

**DOI:** 10.1101/2025.07.04.663236

**Authors:** Layale Youssef, Amanda O’Farrell, Nesrine Harroum, Younes Bakhta, Liora Cohen, Benjamin Pageaux, Jason L. Neva

**Affiliations:** École de kinésiologie et des sciences de l’activité physique (EKSAP), Faculté de médecine, Université de Montréal, Montréal, Québec, Canada; Centre de Recherche de l’Institut Universitaire de Gériatrie de Montréal (CRIUGM), Montréal, Québec, Canada; Centre Interdisciplinaire de Recherche sur le Cerveau et l’Apprentissage (CIRCA), Montréal, Québec, Canada

**Author notes:** Both authors equally contributed to this work (Benjamin Pageaux, Jason L. Neva). ***Corresponding Author*:** Layale Youssef, 4545 Queen Mary Road, Montreal, QC, Canada.

**Keywords:** eccentric cycling, concentric cycling, neuroplasticity, TMS, primary motor cortex

## Abstract

Acute aerobic exercise (AAE) can modulate primary motor cortex (M1) excitability. To date, studies evaluating its effects have focused almost exclusively on concentric cycling. Critically, we found that eccentric AAE enhances motor learning more than concentric AAE, possibly explained by enhanced frontal-parietal brain activation during eccentric cycling. Yet, M1 excitability mechanisms underlying this eccentric AAE-enhanced motor learning remain unknown. Thus, this study aimed to evaluate the effect of eccentric cycling AAE on M1 excitability using transcranial magnetic stimulation (TMS). Thirty adults performed three 20 min-conditions: *i)* eccentric cycling AAE, *ii)* concentric cycling AAE, and *iii)* rest. Cycling AAE was carried out at a workload corresponding to 70% of peak heart rate (%HR_peak_) measured during concentric incremental cycling exercise. TMS assessments were conducted before (Pre), immediately (Post_0_) and 20 minutes after (Post_20_) AAE/rest to evaluate changes in corticospinal excitability (CSE) and short-interval intracortical inhibition (SICI). Overall, we found CSE increased and intracortical inhibition (SICI) was reduced at Post_20_ to a comparable extent following eccentric and concentric cycling AAE compared to rest. Also, %HR_peak_, muscle pain and perceived effort were lower during eccentric cycling AAE compared to concentric cycling AAE. Our results showed that eccentric cycling impacted M1 excitability change to a comparable degree as concentric cycling, while requiring less cardiovascular response, eliciting less muscle pain and lower perceived effort. Taken together, our results suggest that eccentric cycling AAE may be a valuable intervention to modulate M1 excitability for populations with limited cardiovascular capacity and may have potential implications in clinical and sports-related contexts.

## Introduction

Acute aerobic exercise (AAE) has been shown to promote neuroplasticity ^1–4^ and motor learning ^5,6^. Transcranial magnetic stimulation (TMS) studies have demonstrated that a single bout of cycling aerobic exercise can modulate primary motor cortex (M1) excitability and enhance responsiveness to neuroplasticity-inducing protocols^1,4^. This suggests that AAE may serve a preparatory or “priming” role in facilitating neural plasticity. Further work using methods of single- and paired-pulse TMS has shown that lower-limb cycling AAE modulates inhibitory and facilitatory circuits within the non-exercised upper-limb representation in M1^1,3,4^. Our recent meta-analysis showed that short-interval intracortical inhibition (SICI) was most consistently decreased and corticospinal excitability (CSE) can be increased^1^. However, the vast majority of studies assessing M1 excitability changes following lower limb cycling AAE have used the concentric muscle contraction modality, commonly referred to as concentric cycling. As a result, the impact of the eccentric muscle contraction mode during lower limb cycling AAE on M1 excitability remains underexplored.

Eccentric cycling is a less conventional form of lower-limb exercise involving muscle lengthening contractions while resisting motor-driven pedals. Compared with concentric cycling, it allows greater power output with lower cardiovascular, respiratory, and metabolic demands^7^, and has shown promising applicability for individuals with cardiorespiratory limitations ^8^. As a result, there has been growing interest in incorporating eccentric cycling into clinical rehabilitation settings ^8^ as well as in experimental research ^7^.

Eccentric contraction exercise, such as downhill treadmill walking, has been shown to increase CSE in non-exercised upper limb M1 representation to a similar extent as concentric contractions during uphill treadmill walking ^9^. Additionally, eccentric contraction exercise of the upper-limbs induced prolonged changes in intracortical inhibition as compared to concentric contraction exercise ^10^. Recent research demonstrated that eccentric cycling AAE leads to greater activation in motor-related brain areas, like prefrontal and parietal cortices, compared to concentric cycling AAE ^11^. Importantly, the prefrontal cortex plays a key role in cognitive, explicit, and executive processes, which support the early stages of motor learning ^12^. Additionally, modulation of CSE ^13^ and SICI circuitry ^14^ have been associated with motor skill acquisition and learning. Our recent work also showed that cycling AAE can preferentially enhance M1 interneuron excitability as assessed by distinct TMS current directions ^2,15^. Specifically, M1 excitability assessed with anterior-to-posterior TMS current, which preferentially activates unique interneurons associated with task-related prefrontal cortical engagement ^16^ and motor learning ^17^, was enhanced to a greater extent than traditional posterior-to-anterior TMS ^2,15^. Since eccentric cycling exercise is associated with greater prefrontal cortical activation ^11^, it is possible that eccentric AAE may impact M1 interneuron excitability as measured by anterior-to-posterior TMS to a greater extent than concentric AAE. Taken together, these findings suggest that eccentric cycling exercise may engage unique motor-related brain circuits involved in the control of cognitive and motor function, potentially facilitating skill acquisition and motor learning.

We recently found that eccentric cycling AAE enhanced motor skill acquisition and learning more than concentric cycling ^18^. Consistent with previous work ^5,6^, concentric cycling AAE enhances motor learning compared to rest, and our findings suggest that eccentric cycling AAE may promote greater modulation of M1 excitability than concentric cycling AAE, supporting the enhanced effect of eccentric cycling AAE ^18^. Eccentric contractions are known to involve distinct neural control strategies compared to concentric contractions, including greater cortical involvement during movement execution ^19^. Moreover, corticospinal excitability exhibits a specific temporal pattern during eccentric contractions, characterized by a reduction prior to movement followed by an increase after execution ^20^. This distinct neural signature may reflect the greater complexity of eccentric motor control and could contribute to differential modulation of M1 excitability.” To the best of our knowledge, only one study has examined the effects of eccentric cycling on CSE in a non-exercised upper-limb muscle and reported no significant changes following exercise ^21^. However, the study compared eccentric and concentric cycling conditions without including a rest (a non-exercise control) condition. Critically, we address this limitation within the current study design.

Thus, the aim of this study was to assess changes in M1 excitability following eccentric cycling AAE as compared to concentric cycling AAE and to rest. We hypothesized that eccentric cycling AAE will result in greater changes in M1 excitability, specifically a greater increase in CSE and a more pronounced reduction in SICI, compared to concentric cycling AAE. We also expected to find a decrease in SICI following concentric cycling AAE compared to rest. Finally, for all these effects, we expected to find a greater increase in CSE and decrease in SICI as measured with anterior-to-posterior TMS current.

## Methods

### Participants

Thirty healthy adults (28.03 ± 8.88 years, 15 male and 15 female), all right-handed (Edinburgh Handedness Inventory: 93.80 ± 12.36; ^22^), were enrolled in this study. This was done to standardize the tested hemisphere and reduce inter-individual variability, as prior TMS studies have reported hemispheric asymmetries in corticospinal excitability in right-handed individuals ^23^. The characteristics of participants are presented in Table 1. A sensitivity analysis performed in G*Power with an alpha risk of 0.05 and a sample size of 30 indicated that we had a 90% chance of observing a medium effect size of f(U) = 0.272 (∼ *η²_p_* = 0.069) or higher ^24^. The experimental procedures were approved by the Aging and Neuroimaging Research Ethics Board of the Research Center of the Montreal University Geriatric Institute (CRIUGM; CER VN 23-24-37), and written informed consent was obtained from all participants prior to their involvement in the study. All participants were assessed for any potential contraindications to TMS through standard screening forms. They indicated no history of neurological disorders, and the Physical Activity Readiness Questionnaire (PAR-Q) indicated they were all in adequate physical condition to complete the exercise protocols. All participants filled in the International Physical Activity Questionnaire (IPAQ) to estimate their physical activity level.

**Table 1:** Participants’ characteristics.

| Characteristics | N = 30 |
| --- | --- |
| Sex (Men/Women) | 15/15 |
| Age (years) | $28.03 \pm 8.88$ |
| Handedness (EHI score) | $93.83 \pm 12.36$ |
| Weight (kg) | $69.93 \pm 14.06$ |
| IPAQ (METs) | $4526.7 \pm 2832.1$ |
| HR<sub>peak</sub> (bpm) | $174 \pm 13$ |
| Peak power output (W) | $182 \pm 49$ |

Data are presented as mean ± SD. EHI = Edinburgh Handedness Inventory, IPAQ = International physical activity Questionnaire, MET = Metabolic equivalent tasks; HR<sub>peak</sub> = Heart rate peak (measured), bpm = beats per minute, W = Watts.

### Experimental design

Each participant attended three experimental sessions, with at least 4 days between each session. Figure 1 represents an overview of the experimental design. The experimental conditions included: *i)* seated rest, *ii)* concentric cycling exercise (CON), and *iii)* eccentric cycling exercise (ECC) with each lasting approximately 2.5 hours. The first visit involved the resting experimental condition followed by a concentric incremental cycling exercise test to determine the exercise parameters for the subsequent sessions. The incremental exercise test was then followed by eccentric cycling familiarisation. The following two visits consisted of 20 minutes of moderate-intensity exercise, one involving concentric cycling exercise and the other eccentric cycling exercise and were performed in a counterbalanced order (half of the participants underwent concentric cycling exercise before eccentric cycling exercise). Neurophysiological measurements were collected at three time points: before (Pre), immediately after (Post_0_; starting immediately post exercise), and 20 minutes after (Post_20_; starting 20 min post exercise) the experimental intervention. Neurophysiological measurements lasted up to 12 min. To account for potential diurnal variations in CSE, all visits were scheduled at the same time of day (± 2 hours) for each participant ^25^.

**Figure 1:**
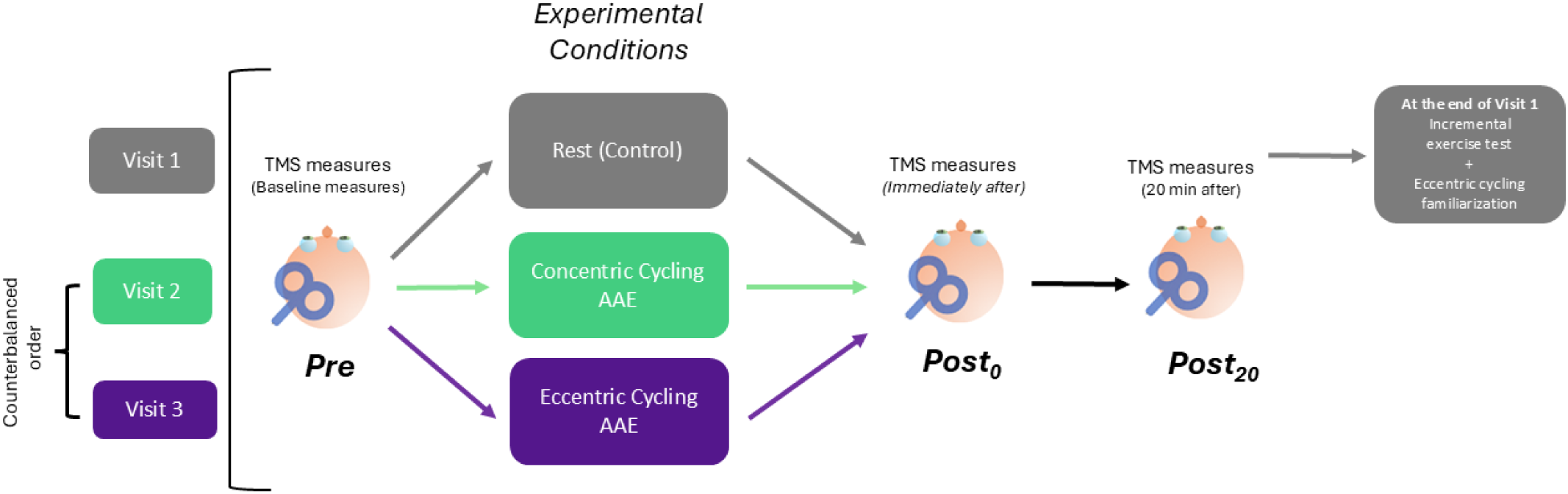
Overview of the study design. Visit 1 consisted of the resting experimental condition followed by an incremental exercise test to determine the exercise parameters for the subsequent sessions. Visits 2 and 3 consisted of the moderate intensity cycling exercise conditions, one involving 20 min of concentric cycling and the other 20 min of eccentric cycling and were performed in a counterbalanced order. Neurophysiological measurements were collected at three time points: Pre (before cycling exercise/rest), Post_0_ (immediately after cycling exercise/rest) and Post_20_ (20 min after cycling exercise/rest). At each time timepoint, MEP 110%, MEP 130% and SICI were obtained. TMS = transcranial magnetic stimulation; AAE = acute aerobic exercise; MEP = motor-evoked potential; SICI = short-interval intracortical inhibition.

### Incremental test

A concentric incremental cycling exercise test was conducted on a concentric recumbent cycle ergometer (CY00500, Cyclus 2, Germany) to determine the exercise intensity for the subsequent visits. The starting workload and incremental increases were tailored to each participant’s body weight (e.g., 65W for 65 kg) ^26^. Participants were instructed to maintain a steady cycling cadence of 60 rpm while seated comfortably on the ergometer. The starting workload, as well as the increments that followed, were scaled to each participant’s body weight and increased every 2 minutes until exhaustion ^26,27^. For example, participants weighing approximately 50 kg started at 50 W with 15 W increments, whereas those around 60 kg began at 60 W with 20 W increments. Proportionally higher starting workloads and increments were applied for heavier participants (see Table S1 in the supplementary material for details). On average, the duration of the incremental exercise test was between 10 and 12 minutes. Exhaustion was defined as the inability to maintain a cadence at 60 rpm for at least 10 seconds, despite verbal encouragement. Heart rate was continuously monitored (Polar H10, Polar Electro 2024, Finland), and peak heart rate (HR_peak_) was recorded at exhaustion. Participants rated their perceived effort and muscle (quadriceps) pain using the CR100 scale ^28,29^ at the end of each 2-minute increment. Additionally, their affect was assessed using the Feeling scale at the end of each increment ^30^.

### Cycling exercise and rest

Before the experimental sessions, participants performed a 20-minute familiarization session on the eccentric cycle ergometer. The power output was adjusted to correspond to 50% of HR_peak_, as established during the incremental test. Participants were instructed to maintain a cadence of 60 rpm while resisting the motor-driven pedals in order to produce eccentric muscle contractions. Visual feedback for cadence and power was displayed throughout the session: the cadence bar turned green when the target cadence was achieved, and the power indicator turned white when the target resistance was reached. The ability to keep the cadence within the green target zone at the end of the 20-minute cycling bout was considered an indication that participants were adequately familiarized with the task.

The cycling AAE intervention lasted 25 minutes, consisting of a 5-minute warm-up followed by 20 minutes of continuous moderate-intensity cycling exercise. The cycling power for each participant was determined based on their HR_peak_, measured during the incremental exercise test, to ensure the intensity of the exercise corresponded to a moderate intensity. During the 5-minute warm-up, participants cycled at a power equivalent to 50% of their HR_peak_ (low intensity), and the subsequent 20-minute continuous cycling exercise was performed at 70% of their HR_peak_ (moderate intensity). These intensity levels were selected based on the American College of Sports Medicine (ACSM) guidelines ^26^. For both concentric and eccentric cycling AAE, the target power output was derived from the workload corresponding to 70% of the participant’s HRpeak during the concentric incremental test. This same absolute power output was then maintained throughout both the concentric and eccentric cycling sessions. Thus, the two exercise conditions were intentionally matched for external workload rather than for the physiological response elicited during exercise, such as HR or oxygen consumption. We selected this approach because eccentric cycling is characterized by lower cardiopulmonary, metabolic, and perceptual demands than concentric cycling when performed at the same absolute power output. Matching conditions based on HR or oxygen consumption would have required substantially higher eccentric power outputs ^7,31^, thereby altering the research question from a comparison at equivalent mechanical workload to a comparison at equivalent internal physiological load. The present approach is consistent with previous eccentric cycling studies using concentric-derived workloads ^7,11^ and preserves the clinically relevant feature of eccentric cycling, which is the ability to perform a given mechanical workload with reduced physiological and perceptual demand.

During both eccentric (Cyclus 2, Germany; model CY00360) and concentric (Cyclus 2, Germany; model CY00500) cycling, the ergometers operated in isopower mode, with participants instructed to maintain a constant cadence of 60 rpm. The concentric and eccentric ergometers shared identical bike frames; specifically, the motor for concentric cycling (CY00500) was mounted on the same frame model used for the eccentric ergometer. Heart rate was monitored continuously and recorded every 5 min during each intervention (Polar H10, Polar Electro Oy, Kempele, Finland). Mean heart rate during each 20-min cycling/rest session was calculated and expressed relative to each participant’s HR_peak_ obtained during the concentric incremental test (%HR_peak_ = mean session HR / HR_peak_ × 100). Additionally, participants reported their perceived effort and muscle pain using the CR100 scale ^28,29^, as well as their affect using the Feeling scale (Hardy & Rejeski, 1989) every 5 minutes. During the cycling exercise, participants were instructed to keep their hands relaxed, resting them gently on top of the handlebars without gripping, to reduce activation of the non-exercised hand muscles. Continuous electromyography (EMG) was recorded from both the right and left abductor pollicis brevis (APB) muscles to confirm that the non-exercised hand muscles remained relaxed.

The rest intervention also lasted 25 minutes, during which participants sat comfortably in a chair and watched an emotionally neutral documentary. They were instructed to keep their upper limbs relaxed throughout the session. Heart rate, perceived effort, muscle pain, and affect were measured every 5 minutes, following the same procedure as the cycling AAE interventions.

### Muscle pain assessments between exercise sessions

After visits 2 and 3, participants were provided with forms to assess their muscle pain at home in three different contexts: “when standing”, “when walking”, and “when taking the stairs up and down”. Participants were asked to rate their pain intensity using a visual analog scale, marking a vertical line on a 10-centimeter line ranging from 0 to 10, where 0 represented “no pain” and 10 represented “maximum pain”. Pain assessments were completed at 24-, 48-, 72-, and 96-h post-exercise.

### Electromyographic recording

During all TMS measurements, EMG recordings were obtained from the right abductor APB muscle. In a belly-tendon configuration, 1 cm diameter electrodes (Covidien, Mansfield, MA, USA) were placed over the APB, with the ground electrode positioned on the ulnar styloid. EMG recordings were obtained using LabChart software (LabChart 8.0), with signals sampled through a PowerLab data acquisition system (PL3516 PowerLab, 16/35 16 Channel Recorder, AD Instruments, Colorado Springs, CO, USA) and a bioamplifier (Dual Bio Amp, AD Instruments, Colorado Springs, CO, USA). The data were acquired at a rate of 2 kHz, with a bandpass filter (20-400 Hz) and a notch filter set to a center frequency of 50 Hz. Data were captured in a 500-ms sweep, starting 100 ms before and ending 400 ms after the delivery of TMS.

### Transcranial magnetic stimulation

Participants were comfortably seated on an adjustable chair and remained at rest during TMS measurements. A Magstim BiStim 200^2^ stimulator (Magstim Co., UK) was used, connected to a 70 mm figure-of-eight coil (Magstim 70 mm P/N 9790, Magstim Co., UK) to deliver monophasic TMS pulses. The TMS current within the coil was adjusted to produce current flow in either a posterior-to-anterior (PA) or anterior-to-posterior (AP) direction. The standard coil delivered TMS pulses in a PA current direction. A custom coil was designed to deliver an AP current, generating a TMS pulse in the opposite direction (i.e., anterior-to-posterior) to the standard PA TMS coil. The TMS coils were positioned at a 45° angle to the mid-sagittal plane, with the handle facing posteriorly. To generate pulses in the lateral-medial (LM) direction, the standard TMS coil was used by rotating the handle 90° to the mid-sagittal plane ^17,32^. To ensure accurate coil positioning and continuous monitoring, Brainsight neuronavigation software (Rogue Research Inc., Montreal, QC, Canada) was used throughout the study using a template brain. The standard TMS coil over M1 to identify the “hotspot” associated with the APB representation, was then utilized for all TMS current directions (PA, AP, and LM). Resting motor threshold (RMT) was then determined at this site for each of the TMS current directions (PA, AP, and LM). For each current direction, TMS pulses were delivered at a constant intensity relative to the corresponding RMT throughout the session. RMT was defined as the minimum intensity required to evoke 5 out of 10 consecutive motor evoked potentials (MEPs) with a peak-to-peak amplitude of at least 50 μV. Throughout the study, TMS pulses were delivered at a random frequency between 0.15 and 0.2 Hz, with approximately 20% variation. TMS measures were collected in the following order: MEPs at 110% and then 130% of RMT, and SICI with PA TMS current; next, MEPs at 150% of RMT with LM TMS current (only collected at Pre); and finally, MEPs at 110% and then 130% of RMT, and SICI with AP TMS current.

### MEP amplitudes

To investigate potential changes in CSE following cycling AAE, MEP amplitudes were measured in both the PA and AP current directions. MEPs were recorded at 110% RMT, as previous research suggests that lower suprathreshold TMS intensities are necessary to preferentially activate distinct interneuron circuits with PA and AP currents ^17,32^. Our previous research demonstrated that moderate-intensity AAE increased CSE, as measured with the lower intensity (i.e., 110% RMT) in the AP current direction but not in the PA direction and had no effect at higher TMS intensities (i.e., 130% RMT and above) ^2^. MEPs were also assessed at 130% RMT to confirm our prior findings following concentric cycling AAE ^2^ and to further examine the effect of eccentric cycling AAE on these measures. A total of fifteen stimuli were applied at each TMS intensity and current direction. The intervals between TMS measurements were typically 10-15 seconds, with transitions between the PA and AP coil orientations taking approximately 1 minute.

### SICI

The effect of cycling AAE on intracortical inhibition in the PA and AP current directions was evaluated using short-interval intracortical inhibition (SICI), following methods established in previous research ^2^. A subthreshold conditioning stimulus (CS) was delivered prior to a suprathreshold test stimulus (TS) at the APB hotspot over M1. The CS was set to 80% RMT, while the TS intensity was adjusted to elicit an average peak-to-peak MEP amplitude of approximately 1 mV across 15 stimulations. For the PA current direction, an interstimulus interval (ISI) of 2 ms was chosen based on prior studies showing a reduction in inhibition following acute exercise ^2,3^. In contrast, a 3 ms ISI was utilized for SICI measurements in the AP current direction, as this interval has been shown to elicit more stable and robust inhibition compared to the 2 ms ISI ^2^. Also, this adjustment accounts for the longer cortical transmission pathways associated with AP TMS ^17^, making the 3 ms ISI a more suitable choice for assessing SICI in this direction. At each time point (Pre, Post_0_, and Post_20_), SICI was measured by delivering 15 paired pulses (CS followed by TS) and 15 single TS for both the PA and AP current directions.

### MEP onset latencies

The earliest onset latency among a set of 15 MEPs was identified to indirectly assess I-wave recruitment and infer the preferential activation of specific interneuron populations. This analysis was performed for both PA and AP TMS current directions, with stimulation set at 110% RMT. To estimate D-wave activation, the same procedure was applied but using an LM TMS current direction at 150% RMT ^32^.

### Data processing and statistical analysis

Repeated measures analysis of variance (RM-ANOVA) was conducted to assess the effects of concentric AAE, eccentric AAE and rest on heart rate and perceptual responses (perceived effort, muscle pain, and affect) and neurophysiological measures obtained using TMS (MEP amplitude, SICI, and MEP onset latency). Specific analyses are detailed below. Where appropriate, post hoc comparisons were performed and adjusted for multiple comparisons with the Holm-Bonferroni correction. Significance was set at a *p* < .05. Effect sizes were calculated and expressed as partial eta squared (η²_p_) for ANOVAs (with 0.01, 0.06, and 0.14 indicating small, moderate, and large effects, respectively) and as Cohen’s d for pairwise comparisons (with 0.2, 0.5, and 0.8 indicating small, moderate, and large effects, respectively). All statistical analyses were conducted in R software (4.2.1, foundation for statistical computing, Vienna, Austria).

### Cycling AAE / rest data

To determine whether the AAE or rest conditions (ECC, CON, REST) elicited distinct physiological and psychological responses, we analyzed %HR_peak_, perceived effort, muscle pain, and affect data during the sessions. Two separate 2-way RM-ANOVAs were conducted. The first analysis examined average heart rate peak and affect, with CONDITION (ECC, CON, REST) and TIME (5, 10, 15, and 20 min) as within-subject factors. In the second analysis, REST was excluded for perceived effort and muscle pain, as all participants reported a score of zero throughout the intervention, allowing for a direct comparison between ECC and CON.

For the analysis related to the pain assessments after the cycling AAE interventions, 2-way RM- ANOVAs were conducted for each assessment with CONDITION (ECC, CON) and TIME (24h, 48h, 72h and 96h) as within-subject factors.

### Neurophysiological data

#### RMT, TS % maximum stimulator output (MSO) and TS MEP amplitudes

To ensure consistency in RMT (%MSO) at the start of each session (pre-AAE/rest), a two-way RM-ANOVA was performed with CONDITION (ECC, CON, REST) and TMS CURRENT (PA, AP) as within-subject factors. The stability of TS %MSO values during SICI assessment before and after cycling AAE/rest was verified using a three-way RM-ANOVA, incorporating TIME (Pre, Post_0_, Post_20_), CONDITION (ECC, CON, REST), and TMS CURRENT (PA, AP) as within-subject factors. This confirmed that TMS intensity remained stable throughout SICI assessments across conditions, time and current directions. Similarly, a three-way RM-ANOVA was conducted to assess the stability of TS MEP amplitudes before and after cycling AAE/rest, considering TIME (Pre, Post_0_, Post_20_), CONDITION (ECC, CON, REST), and TMS CURRENT (PA, AP). This analysis verified that CSE remained consistent across the different AAE conditions, across time and TMS currents during SICI assessment.

#### MEP onset latency

A semi-automated system was utilized to calculate MEP onset latency, defined as the moment when the rectified EMG signal exceeded five times the average pre-stimulus EMG level. MEP latencies induced by single-pulse TMS were determined for each current direction (PA, AP, LM). Consistent with prior studies ^2,17,32^, differences in MEP latencies (ΔPA-LM, ΔAP-LM) were employed as indirect markers of I-wave recruitment, providing evidence for the activation of specific interneuron circuits. To analyze the earliest MEP latency responses, we conducted a two-way RM-ANOVA with TMS CURRENT (PA, AP, LM) and CONDITION (ECC, CON, REST) as within-subject factors. In a separate analysis, differences in MEP onset latency between PA-LM and AP-LM were assessed using another two-way RM-ANOVA, with MEP ONSET DIFFERENCE (ΔPA-LM, ΔAP-LM) and CONDITION (ECC, CON, REST) as within-subject factors.

#### EMG and MEP data processing for MEP amplitudes and SICI

For both MEPs (collected at 110% and 130% RMT) and SICI, the EMG data were carefully examined for any voluntary muscle activity. Trials with any voluntary pre-stimulus EMG activity were excluded from analysis, accounting for 1.2% of the total trials. Peak-to-peak MEP amplitudes (in millivolts, mV) were analyzed using custom MATLAB scripts. SICI was quantified as the ratio of CS + TS amplitude to TS MEP amplitude: SICI ratio = (CS + TS) / TS. In this ratio, lower values reflect stronger inhibition, while higher values indicate reduced inhibition (or disinhibition). For clarity, throughout the manuscript, the term “SICI” refers to the physiological construct of intracortical inhibition, whereas “SICI ratio” refers to its operational quantification (i.e., SICI ratio = (CS + TS)/TS). Hence, changes in the SICI ratio are interpreted inversely with respect to inhibition (i.e., a decrease in SICI ratio value indicates increased intracortical inhibition, whereas an increase in SICI ratio value reflects reduced intracortical inhibition).

#### MEP amplitudes

To evaluate the effect of eccentric and concentric cycling AAE on CSE, mean MEP amplitude was calculated from fifteen responses at each of the two TMS intensities (110% and 130% RMT). Each intensity level was analyzed separately, as lower stimulation intensities (e.g., 110% RMT) increase the likelihood of preferentially engaging distinct interneuron circuits depending on the TMS current direction (PA or AP) ^16,17^. To examine these effects, three-way RM-ANOVAs were conducted for each TMS intensity (110% and 130% RMT), with TIME (Pre, Post_0_, Post_20_), CONDITION (ECC, CON, REST), and TMS CURRENT (PA, AP) as within-subject factors, using mean MEP amplitude as the dependent variable.

#### Short-interval intracortical inhibition

To evaluate the effect of eccentric and concentric cycling AAE on SICI, the mean MEP amplitude was calculated for each of the fifteen TS and CS+TS pulses for both TMS current directions (PA, AP) to determine the SICI ratio. A three-way RM-ANOVA was then performed with TIME (Pre, Post_0_, Post_20_), CONDITION (ECC, CON, REST), and TMS CURRENT (PA, AP) as within-subject factors, using the SICI ratio as the dependent variable.

## Results

### Baseline physical activity level, cycling exercise, and rest data

Based on the IPAQ classification, 10 participants were categorized as having low activity, 12 as moderate activity, and 8 as high activity. Heart rate, perceived effort, muscle pain, and affect were recorded every 5 minutes during the 25-minute experimental interventions. Figure S1 presents the mean values for each intervention, while Figure S2 illustrates their time course.

*% HR_peak_*. Figure S1A. There was a main effect of CONDITION [*F_2,58_* = 183.924, *p* < .001, *η²_p_ =* 0.864]. Post hoc analyses revealed that % HR_peak_ was higher for the concentric cycling AAE compared to both eccentric cycling AAE (*p* < .001, *d* = 1.039) and rest (*p* < .001, *d* = 2.306), and eccentric cycling AAE showed a higher % HR_peak_ than rest (*p* < .001, *d* = 1.019). There was no main effect of TIME [*F_3,87_* = 0.424, *p* = .736, *η²_p_ =* 0.014], and no CONDITION x TIME interaction [*F_6,174_* = 0.185, *p* = .981, *η²_p_ =* 0.006].

### Affect

Figure S1B. There was a main effect of CONDITION [*F_2,58_* = 36.498, *p* < .001, *η²_p_ =* 0.560]. Post hoc analysis revealed that affect was higher at rest compared to both concentric (*p* < .001, = 1.134) and eccentric (*p* < .001, *d* = 0.602) cycling AAE, and eccentric cycling AAE showed a higher affect compared to concentric cycling AAE (*p* <.001, *d* = 0.654). There was no main effect of TIME [*F_3,87_* = 0.354, *p* = .787, *η²_p_ =* 0.012] and no CONDITION x TIME interaction [*F_6,174_* = .102, *p* = .996, *η²_p_ =* 0.004].

#### Perceived effort

Figure S1C. There was a main effect of CONDITION [*F_1,29_* = 40.083, *p* < .001, *η²_p_ =* 0.580] revealing that perceived effort was higher during concentric cycling AAE compared to eccentric cycling AAE. There was no main effect of TIME [*F_3,87_* = 1.811, *p* = .146, *η²_p_ =* 0.059], and no CONDITION x TIME interaction [*F_3,87_* = 1.351, *p* =.259, *η²_p_ =* 0.045].

#### Muscle pain

Figure S1D. There was a main effect of CONDITION [*F_1,29_* = 16.290, *p* < .001, *η²_p_ =* 0.360] revealing that muscle pain was higher during concentric cycling AAE compared to eccentric cycling AAE. There was no main effect of TIME [*F_3,87_* = 1.019, *p* = .388, *η²_p_ =* 0.034], and no CONDITION x TIME interaction [*F_3,87_* = 0.135, *p* = .939, *η²_p_ =* 0.005].

### Muscle pain post-AAE

Muscle pain *i)* when standing, *ii)* when walking and *iii)* when taking the stairs up and down, were evaluated 24h, 48h, 72h, and 96h post-AAE (Figure S3).

#### Muscle pain when standing

Figure S3A. There was a CONDITION x TIME interaction (F_3,87_ = 26.593, *p <* .001, *η²_p_* = 0.478). Post-hoc analyses revealed higher pain following 24h (*p* < .001, *d* = 1.610), 48h (*p* < .001, *d* = 1.808) and 72h (*p* = .014, *d* = 0.876) following eccentric cycling AAE compared to concentric cycling AAE. Also, following eccentric cycling AAE, pain decreased from 48h to 72h (*p* < .001, *d* = 1.368) and from 72h to 96h (*p* = .024, *d* = 0.808). Additionally, there was a main effect of CONDITION (F_1,29_ = 42.206, *p <* .001, *η²_p_* = 0.593) and a main effect of TIME (F_3,87_ = 32.420, *p <* .001, *η²_p_* = 0.528).

#### Muscle pain when walking

Figure S3D. There was a CONDITION x TIME interaction (F_3,87_ = 31.606, *p* < .001, *η²_p_* = 0.522). Post-hoc analyses revealed higher pain following 24h (*p* < .001, *d* = 1.704), 48h (*p* < .001, *d* = 2.234) and 72h (*p* = .003, *d* = 1.010) following eccentric cycling AAE compared to concentric cycling AAE. Also, following eccentric cycling AAE, pain decreased from 48h to 72h (*p* < .001, *d* = 1.778) and from 72h to 96h (*p* = .002, *d* = 1.069). Additionally, there was a main effect of CONDITION (F_1,29_ = 62.836, *p <* .001, *η²_p_* = 0.684) and a main effect of TIME (F_3,87_ = 46.254, *p <* .001, *η²_p_* = 0.615).

### Muscle pain when taking the stairs up and down

Figure S3G. There was a CONDITION x TIME interaction (F_3,87_ = 24.502, *p* < .001, *η²_p_* = 0.458). Post-hoc analyses revealed higher pain following 24h (*p* < .001, *d* = 1.608), 48h (*p* < .001, *d* = 1.778) and 72h (*p* = .013, *d* = 0.868) following eccentric cycling AAE compared to concentric cycling AAE. Also, following eccentric cycling AAE, pain decreased from 48h to 72h (*p* < .001, *d* = 1.835) and from 72h to 96h (*p* = .008, *d* = 0.928). Additionally, there was a main effect of CONDITION (F_1,29_ = 39.965, *p <* .001, *η²_p_* = 0.579) and a main effect of TIME (F_3,87_ = 47.362, *p <* .001, *η²_p_* = 0.620).

### Neurophysiological data

#### Baseline neurophysiological data

##### RMT values

Table S2. There was a main effect of CURRENT (F_2,58_ = 69.368, *p* < .001, *η²_p_* = 0.705). Post hoc analyses revealed that RMT for PA TMS current was lower than AP TMS current (*p* < .001, *d* = -2.564), RMT for LM TMS current was lower than AP TMS current (*p* < .001, *d* = 1.138), and RMT for PA TMS current was lower than LM TMS current (*p* < .001, *d* = 0.634). Moreover, there was no main effect of CONDITION (F_2,58_ = 0.482, *p* = .620, *η²_p_* = 0.016) or CURRENT x CONDITION interaction (F_4,116_ = 1.434, *p* = .227, *η²_p_* = 0.047).

##### TS% MSO

Table S3. There was a main effect of CURRENT (F_1,29_ = 104.537, *p* < .001, *η²_p_* = 0.783) revealing higher values for AP TMS current compared to PA TMS current. Additionally, there was no main effect of CONDITION (F_2,58_ = 0.622, *p* = .540, *η²_p_* = 0.021) or TIME (F_2,58_ = 2.812, *p* =.068, *η²_p_* = 0.088), nor interactions (all *ps* > .913).

##### TS MEP amplitudes

Table S3. TS MEP amplitudes during SICI were consistent across time, between exercise conditions and between TMS currents. More specifically, there were no main effects of CONDITION (F_2,58_ = 1.038, *p* = .360, *η²_p_* = 0.035), TIME (F_2,58_ = 2.565 *p* = .085, *η²_p_* = 0.081) or CURRENT (F_1,29_ = 2.428 *p* = .130, *η²_p_* = 0.077), nor interactions (all *ps* > .774).

#### MEP onset latency

Figure S4B. There was a main effect of CURRENT (F_2,58_ = 278.360, *p* < .001, *η²_p_* = 0.906) with post hoc analyses showing a shorter MEP onset latency using LM current compared to PA (*p* < .001, *d* = -2.352) and AP (*p* < .001, *d* = -3.413) TMS current. Additionally, MEP onset latency using PA TMS current was shorter compared to AP TMS current (*p* < .001, *d* = -1.881). Moreover, there was no main effect of CONDITION (F_2,58_ = 0.208 *p* = .812, *η²_p_* = 0.007) or CONDITION x CURRENT interaction (F_4,116_ = 1.113, *p* = .354, *η²_p_* = 0.037).

#### MEP onset latency difference

Figure S4C. There was a main effect of MEP ONSET DIFFERENCE (F_1,29_ = 226.285, *p* < .001, *η²_p_* = 0.886) revealing a greater ΔAP-LM compared to ΔPA- LM. Additionally, there was no main effect of CONDITION (F_2,58_ = 1.537 *p* = .224, *η²_p_* = 0.050) or MEP ONSET DIFFERENCE x CONDITION interaction (F_2,58_ = 0.906, *p* = .411, *η²_p_* = 0.030).

### Baseline CSE and SICI measures

To ensure that CSE and SICI (expressed as SICI ratio) did not differ between the three experimental conditions prior to the intervention, a two-way repeated-measures ANOVA was conducted. *Baseline CSE.* For MEP 110%, there were no main effects of CONDITION (F_2,58_ = 0.231, *p* = .794, *η²_p_* = 0.008), CURRENT (F_1,29_ = 0.095, *p* = .760, *η²_p_* = 0.003) or CONDITION x CURRENT interaction (F_2,58_ = 0.208, *p* = .813, *η²_p_* = 0.007). For MEP 130%, there were no main effects of CONDITION (F_2,58_ = 0.138, *p* = .871, *η²_p_* = 0.005), CURRENT (F_1,29_ = 0.642, *p* = .429, *η²_p_* = 0.022), or CONDITION x CURRENT interaction (F_2,58_ = 0.016, *p* = .984, *η²_p_* = 0.001).

#### Baseline SICI

There was a main effect of CURRENT (F_1,29_ = 13.508, *p* <.001, *η²_p_* = .318) revealing greater SICI ratio values for PA TMS current compared to AP TMS current (i.e. less inhibition for PA TMS current). There were no main effects of CONDITION (F_2,58_ = 0.235, *p* = .791, *η²_p_* = 0.008), or CONDITION x CURRENT interaction (F_2,58_ = 0.060, *p* = .941, *η²_p_* = 0.002).

### Corticospinal excitability

MEP data are displayed for each TMS intensity (110% RMT and 130% RMT) with TMS currents collapsed and separated into PA and AP (Figure 2 and Figure S5 in Supplementary material). Representative individual MEP traces for MEPs at 130% RMT are presented in Figure S6 in Supplementary material.

**Figure 2:**
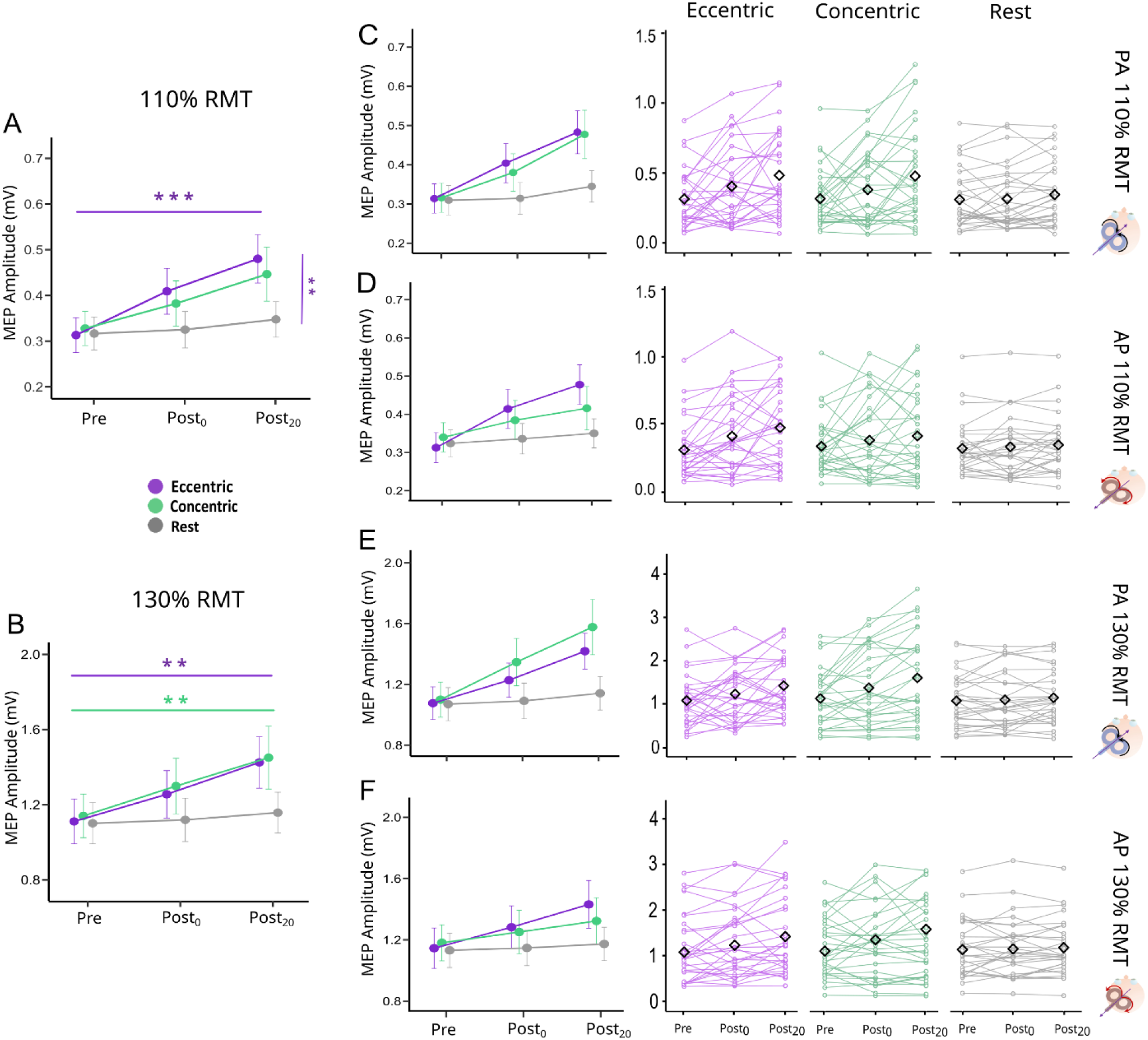
Corticospinal excitability results. Panels A and B display mean peak-to-peak amplitudes with TMS currents collapsed et each time point (110% RMT and 130% RMT respectively). Mean peak-to peak amplitudes with PA at 110% RMT (Panel C), AP at 110% RMT (Panel D), PA at 130% RMT (Panel E) and AP at 130% RMT (Panel F) are displayed at each time point. Plots with individual lines represent individual data corresponding to each previously mentioned panel. Bars represent the standard error of the mean. Purple color refers to eccentric condition, green color refers to concentric condition and grey color refers to rest condition. TMS = transcranial magnetic stimulation. AP = anterior-to- posterior; PA = posterior-to-anterior; MEP: motor-evoked potential; mV: millivolt; RMT = resting motor threshold; Pre: before the experimental intervention; Post_0_: immediately after the experimental intervention; Post_20_: 20 minutes after the experimental intervention. Horizontal purple and green bars represent the difference between Pre and Post_20_ for eccentric and concentric cycling respectively. The vertical purple line represents the difference between eccentric cycling and rest at Post_20_. **: *p-value* < .01; ***: *p-value* < .001.

#### MEPs at 110% RMT

Figure 2A. There was a CONDITION x TIME interaction (F_4,116_ = 3.162, *p* = .016, *η²_p_* = 0.098). Post-hoc analyses revealed a greater MEP amplitude for eccentric cycling AAE at Post_20_ compared to Pre (*p* <.001, *d* = -1.579). At Post_20_, MEP amplitude was greater for eccentric cycling AAE compared to rest (*p* = .002, *d* = 1.121). No significant differences were observed between eccentric and concentric cycling AAE at any time point (all *ps* > .529). Additionally, there were main effects of TIME (F_2,58_ = 19.658, *p* < .001, *η²_p_* = 0.404), and CONDITION (F_2,58_ = 3.398, *p* = .040, *η²_p_* = 0.105) with no other effects or interactions (all *ps* > .975).

#### MEPs at 130% RMT

Figure 2B. There was a CONDITION x TIME interaction (F_4,116_ = 3.027, *p* = .020, *η²_p_* = 0.095). Post-hoc analyses revealed greater MEP amplitude at Post_20_ compared to Pre for both eccentric (*p* = .002, *d* = -1.114) and concentric cycling AAE (*p* = .004, *d* = -1.051). No significant differences were observed between eccentric and concentric cycling AAE at any time point (all *ps* > .838). Additionally, there was a main effect of TIME (F_2,58_ = 21.208, *p* < .001, *η²_p_* = 0.422) and no other effects or interactions (all *ps* > .981).

### Short-interval intracortical inhibition

SICI ratio data are displayed with TMS currents collapsed and separated into PA and AP currents (Figure 3 and Figure S5 in Supplementary material). There was a CONDITION x TIME interaction (F_4,116_ = 2.542, *p* = .043, *η²_p_* = 0.081). Post-hoc analyses revealed higher SICI ratio (reduced inhibition) at Post_20_ compared to Pre for both eccentric (*p* = .004, *d* = -1.060) and concentric cycling AAE (*p* = .047, *d* = - 0.789). At Post_20_, SICI ratio was higher (reduced inhibition) for eccentric cycling AAE compared to rest (*p* = .009, *d* = 0.959). No significant differences were observed between eccentric and concentric cycling AAE at any time point (all *ps* > .197). Additionally, there were main effects of CONDITION (F_2,58_ = 6.546, *p* = .003, *η²_p_* = 0.184), TIME (F_2,58_ = 12.979, *p* <.001, *η²_p_* = 0.309) and CURRENT (F_1,29_ = 6.507, *p* = .016, *η²_p_* = 0.183) with no other interactions (all *ps* > .793).

**Figure 3:**
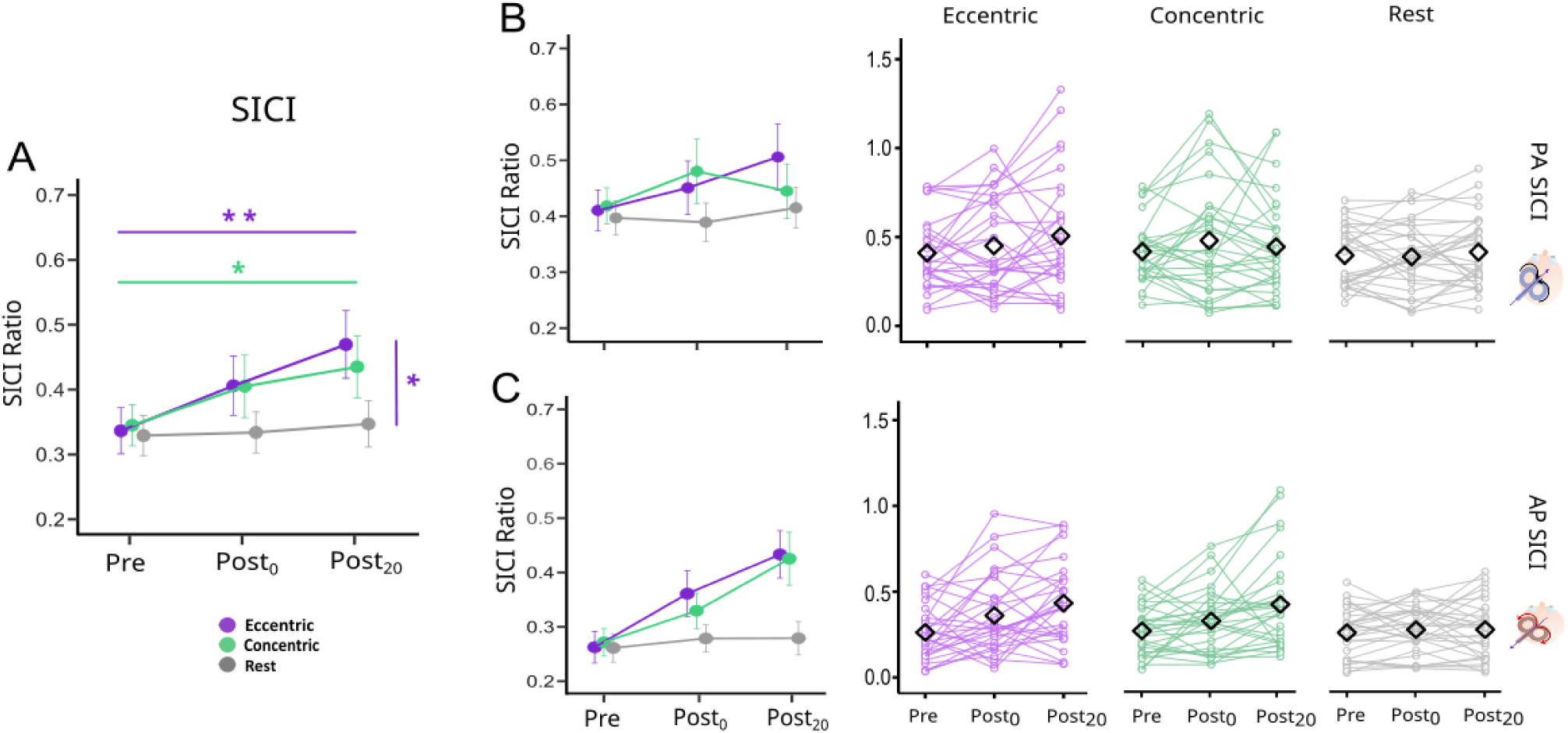
Short-interval intracortical inhibition results. Panel A displays mean SICI ratios with TMS currents collapsed et each time point. Mean SICI ratios with PA (Panel B), and AP (Panel C) are displayed at each time point. Plots with individual lines represent individual data corresponding to each previously mentioned panel. Bars represent the standard error of the mean. Higher values represent less GABAergic inhibition. Purple color refers to eccentric condition, green color refers to concentric condition and grey color refers to rest condition. TMS = transcranial magnetic stimulation. AP = anterior-to-posterior; PA = posterior-to-anterior; MEP: motor-evoked potential; SICI = short-interval intracortical inhibition; Pre: before the experimental intervention; Post_0_: immediately after the experimental intervention; Post_20_: 20 minutes after the experimental intervention. Horizontal purple and green bars represent the difference between Pre and Post_20_ for eccentric and concentric cycling respectively. The vertical purple line represents the difference between eccentric cycling and rest at Post_20_. *: *p-value* < 0.05; **: *p-value* < .01.

## Discussion

In this study, we investigated the impact of an acute bout of eccentric and concentric cycling exercise on M1 excitability. We hypothesized that eccentric cycling AAE would increase CSE and reduce intracortical inhibition (SICI) more than concentric cycling AAE. Our findings partially confirmed our hypothesis: eccentric cycling AAE increased CSE at a lower stimulus intensity (110% RMT), whereas concentric cycling did not. Contrary to our hypothesis, CSE increased similarly following eccentric and concentric cycling AAE at a higher stimulus intensity (130% RMT). Unexpectedly, SICI decreased similarly following both eccentric and concentric cycling AAE. Contrary to our hypothesis, there were no differences in CSE and SICI change across TMS current directions (AP, PA). Finally, eccentric cycling AAE elicited unique psychophysiological responses compared to concentric cycling AAE, with lower %HR_peak_, perceived effort, muscle pain and more positive affective response. Given the comparably similar impact on M1 excitability alongside distinct psychophysiological responses, our findings may have important implications in both clinical and sports-related contexts.

### Acute eccentric and concentric cycling exercise increased corticospinal excitability

It is important to emphasize that all TMS measurements were performed at rest (pre- and post-exercise) and therefore reflect changes in resting corticospinal excitability rather than excitability during active cycling. In addition, because MEP amplitude reflects net corticospinal output, changes in CSE cannot be attributed exclusively to cortical mechanisms and may include cortical, subcortical, and spinal contributions. At low TMS intensity (110% RMT), only eccentric cycling AAE produced a significant increase in CSE. Notably, this increase occurred at the Post_20_ time point, with eccentric cycling AAE showing higher excitability than rest. This increase suggests that eccentric cycling AAE induces time- sensitive neurophysiological changes in M1 excitability. In their study, Fang and colleagues demonstrated that eccentric contractions were associated with greater movement-related cortical potential amplitudes compared to concentric contractions, particularly over sensorimotor areas such as M1, supplementary motor area, and primary somatosensory cortex ^33^.

At higher stimulus intensity (130% RMT), our findings revealed that CSE was enhanced following both concentric and eccentric cycling AAE. Distinct mechanisms may underlie this modulation, yet both cycling AAE types appear to share common results. While our recent meta-analysis suggested that concentric cycling AAE primarily enhances CSE at high intensity AAE, the current study that used moderate intensity also showed an increase. This result aligns with previous studies showing that moderate-intensity concentric cycling AAE may also influence CSE^15^. Hence, our study adds to the growing body of evidence that cycling AAE can enhance M1 excitability ^1^. Interestingly, CSE at 130% RMT was also enhanced 20 minutes following eccentric cycling AAE. A previous study also demonstrated that eccentric muscle contractions required a longer time for movement preparation and a greater magnitude of cortical activity for movement execution compared to concentric contractions ^19^. These findings suggest a heightened cortical activity requirement with eccentric contractions, which is further supported by increased activation in the bilateral prefrontal and right parietal cortex during eccentric cycling ^11^. Importantly, CSE shows a distinct temporal modulation associated with the preparation and execution of eccentric contractions: it tends to decrease before eccentric contraction but increases significantly after execution, displaying a pattern different from that of concentric contractions^20^. The mechanisms underlying these effects remain unclear. Beyond SICI, other intracortical circuits may contribute, as Latella et al. reported selective modulation of long-interval intracortical inhibition (LICI) following eccentric biceps contractions ^10^. Broader brain networks may also be involved, given that eccentric cycling elicits greater frontoparietal activation than concentric cycling and that prefrontal regions can influence M1 excitability ^11^. Finally, spinal or subcortical mechanisms cannot be excluded, as MEP amplitude reflects net corticospinal output. Future studies should assess these mechanisms directly.

It is important to distinguish the present findings from studies assessing CSE during active eccentric contractions or eccentric cycling. For example, Clos et al. (2022) reported lower CSE during eccentric compared with concentric cycling when MEPs were recorded from the exercised knee extensors during active cycling ^34^. Similarly, Duchateau and Enoka (2016) summarized evidence that corticospinal and spinal excitability are reduced during active lengthening contractions, with spinal inhibitory mechanisms largely contributing to this effect ^35^. Yet, our findings reflect resting post-exercise CSE in a non-exercised upper-limb muscle.

In contrast to 110% RMT, M1 excitability changes at 130% RMT were not statistically different from rest at Post_20_ for both concentric and eccentric cycling AAE, which is likely due to greater inter-individual variability at this higher stimulation intensity. MEP amplitudes elicited at 130% RMT are known to be more variable ^36^ which may have limited the ability to detect differences between cycling AAE and rest despite apparent trends. Notably, the individual data at 130% RMT for both cycling AAE conditions at Post_20_ display heterogeneous patterns of excitability change. However, these interpretations remain speculative, and the present findings do not allow us to determine the specific mechanisms underlying the observed changes in CSE following eccentric and concentric cycling compared to non-cycling control conditions (e.g., seated rest). Further work is required to clarify these mechanisms.

### Acute eccentric and concentric cycling exercise decreased intracortical inhibition

Consistent with previous findings, we observed reduced SICI (i.e., increased SICI ratio, indicating reduced intracortical inhibition) following 20 minutes of concentric cycling AAE, aligning with results from our meta-analysis showing a consistent reduction in SICI up to 30 minutes after moderate to high-intensity concentric cycling AAE. This decrease in SICI has been linked to several mechanisms, including GABAergic disinhibition mediated specifically by GABA_A_ receptors ^37^. Notably, our study extends this understanding by demonstrating that eccentric cycling AAE similarly decreases SICI, highlighting that both concentric and eccentric cycling modalities can modulate intracortical inhibition in a relatively similar manner.

The mechanisms underlying this modulation likely involve both shared and distinct pathways. For concentric cycling, the greater cardiorespiratory demand, reflected through a higher heart rate response compared to eccentric cycling, likely increases cerebral blood flow ^38^. This increased blood flow may facilitate the delivery and release of neurochemical mediators such as BDNF, dopamine, and other molecules known to influence M1 excitability and intracortical inhibition ^39,40^. For example, BDNF has been shown to reduce GABA_A_ receptor activity following exercise, potentially contributing to the observed decrease in SICI ^39^. Dopamine, particularly through D2 receptor activation, is also known to decrease GABAergic inhibition and increase cortical excitability ^40^.

In contrast, eccentric cycling AAE induced a lower heart rate response and may not elicit the same increase in cerebral blood flow as concentric cycling AAE. Therefore, additional alternative mechanisms are needed to explain the similar magnitude of SICI reduction observed following eccentric cycling AAE. One possible explanation involves the interaction between cortical inhibitory circuits. Previous research has shown that, compared with concentric contractions, eccentric contractions produce a greater and more prolonged reduction in intracortical inhibition as measured by LICI ^10^. Since LICI has been shown to suppress SICI ^41^, it is plausible that these inhibitory mechanisms interact differently in response to eccentric and concentric cycling AAE. This interaction could account for the reduced intracortical inhibition observed following eccentric cycling compared to rest in the present study. Another potential explanation relates to the different neuromuscular demands of eccentric contractions, which can induce delayed muscle pain despite lower metabolic costs. Previous research has demonstrated that eccentric contractions lead to a decrease in SICI hours after exercise, possibly reflecting widespread effects of delayed muscle pain on M1 excitability ^42^. Moreover, eccentric exercise is associated with an inflammatory response characterized by the secretion of cytokines such as interleukin-6 (IL-6) as compared to concentric exercise ^43^. These cytokines are increasingly recognized as modulators of brain plasticity and may regulate GABA receptor function ^44^. Such neuroimmune interactions could therefore contribute to the decreased SICI observed following eccentric cycling AAE compared to rest in our study. Supporting this notion, participants reported higher muscle pain levels at 24- and 48-hours post-exercise, likely due to eccentric cycling AAE. However, further research is necessary to understand the mechanisms underlying the decreased intracortical inhibition following eccentric cycling AAE.

### Absence of modulation specific to TMS current direction

In this study, we found no significant modulation specific to TMS current direction across CSE and intracortical inhibition measures. Since PA and AP currents preferentially recruit early and later interneuronal circuits reflective of early and late I-waves, respectively ^17,32^, the absence of modulation suggests that neither eccentric nor concentric cycling selectively modulated pathways generating specific I-waves. Although slight changes are observed when inspecting the PA and AP TMS current data separately, the differences did not rise to statistical significance, as our previous findings ^2,15^. Variations in study design (e.g., direct comparison of exercise vs rest) and AAE parameters (e.g., interval vs continuous) compared to the current study may have contributed to a lack of significant difference in M1 excitability measured with PA and AP TMS currents. Since M1 excitability responses measured with AP TMS are known to be more variable and susceptible to inhibition ^17,32^, future work will need to investigate eccentric AAE-induced M1 excitability responses with larger and more diverse sample sizes. Overall, our findings indicate that eccentric cycling and concentric cycling did not selectively alter interneuron circuit excitability mediated by specific I-waves, but rather induced broader changes in CSE and intracortical inhibition.

### Limitations

Our study has limitations. First, eccentric and concentric cycling were matched for external power output, but not for internal physiological demand, such as oxygen consumption, heart rate, or heart rate reserve. The target workload was derived from the concentric incremental test; therefore, a separate eccentric incremental test was not performed. While this design allowed comparison at the same mechanical workload, it cannot determine whether similar neurophysiological responses would occur under conditions matched for physiological demand. Such matching would likely require substantially higher eccentric power outputs ^7,31^, given the lower cardiorespiratory cost of eccentric cycling and the higher peak power outputs reported during eccentric incremental testing. Future studies should directly compare exercise protocols that are matched by workload and physiological demand to determine their influence on exercise-induced changes in M1 excitability. Second, exercise intensity was prescribed based on each participant’s measured HR_peak_. While this is a commonly used method, it may be less precise and more variable than alternative approaches such as oxygen uptake or heart rate reserve. In addition, the HR_peak_ achieved during the incremental test may have been underestimated in some participants, suggesting that maximal cardiorespiratory response may not have been attained; future studies should incorporate gas exchange measures to better verify this. Third, prior research has highlighted the importance of precise timing in detecting transient changes in M1 excitability (Ridding & Ziemann, 2010). In our study, TMS assessments at Pre, Post_0_, and Post_20_ spanned approximately 10– 20 minutes, which may be considered a relatively long testing window. Since exercise-induced modulation of M1 excitability is time-sensitive ^1,2,15^ some short-term effects may have been missed. To better capture these transient changes, future studies might consider limiting the number of post-exercise TMS measurements or reducing the overall assessment duration. Fourth, as spinal excitability was not directly assessed (e.g., via H-reflex measurements), a potential contribution of spinal mechanisms to the observed changes in corticospinal excitability, particularly following eccentric exercise, cannot be fully excluded. Because eccentric contractions induces specific spinal adaptations compared to concentric contractions ^35,45^, future studies should directly assess spinal and subcortical excitability to better distinguish the mechanisms contributing to acute eccentric cycling-induced changes in corticospinal excitability. Finally, TMS outcomes were obtained at rest from APB; therefore, the present findings reflect post-exercise modulation of a non-exercised upper-limb M1 representation and should not be interpreted as changes occurring during cycling or within the exercised lower-limb representation. This choice was aligned with our research question and minimized confounds associated with the exercised muscles, such as exercise-induced neuromuscular fatigue. Future studies should assess both resting and active-state TMS outcomes in exercised and non-exercised muscle representations to determine whether eccentric and concentric cycling differentially influence task-specific and non-task-specific M1 excitability.

## Conclusion

Our study revealed that both concentric and eccentric cycling AAE induced comparable overall changes in M1 excitability, as evidenced by increased CSE and reduced intracortical inhibition. Notable nuances were present in these effects. Eccentric cycling AAE elicited significantly increased CSE at lower stimulation intensity (i.e., 110% RMT), whereas the average increase observed with concentric cycling exercise did not reach statistical significance. At higher stimulation intensities (i.e., 130% RMT), similar increases in CSE were observed following eccentric and concentric exercise. Intracortical inhibition decreased following both exercise modalities, yet only eccentric cycling resulted in reduced inhibition compared to rest 20-min post exercise. Our findings indicate that, regardless of contraction type, an acute bout of cycling exercise can induce neurophysiological changes in M1, which may underpin exercise-enhanced neuroplasticity and motor learning. Importantly, eccentric cycling elicited these effects with lower cardiovascular and perceptual responses than concentric cycling, positioning it as a promising option for individuals with limited cardiovascular capacity. Our findings pave the way for further exploration of the effects of eccentric cycling exercise on M1 excitability, which may lead to tailored exercise strategies applicable in both clinical and sports-related contexts.

## Perspective

Our findings suggest that eccentric and concentric AAE can modulate M1 excitability, supporting the potential of cycling exercise as a priming strategy for motor learning ^18^ and neuroplasticity, with possible clinical applications. These findings also highlight important avenues for future research, including more comprehensive investigation of the M1 circuits and brain regions impacted by eccentric cycling AAE. Assessing other M1 circuits (e.g., LICI) may provide further insight, as prior work suggests these pathways may be differentially modulated following eccentric contractions ^46^. Future studies should also examine the contribution of prefrontal and frontoparietal regions and their interactions with M1 ^11^. Extending this work to different populations and task contexts, including clinical cohorts and studies incorporating direct measures of motor learning, will be essential to better understand how eccentric AAE can be leveraged to enhance neuroplasticity and functional recovery.

From a clinical perspective, eccentric cycling AAE may be particularly relevant, as it can modulate M1 excitability to a comparable degree as concentric cycling AAE while minimizing cardiorespiratory responses and perceived effort, when performed at similar power output than concentric AAE. Incorporating eccentric cycling into rehabilitation programs may provide a lower-demand physiological stimulus to promote M1 excitability, plasticity, and motor learning in older adults with reduced cardiovascular capacity and in patients recovering from stroke.

## Author contributions

**Layale Youssef**: Conceptualization, Methodology, Formal analysis, Investigation, Data curation, Writing - Original draft, Visualization. **Amanda O’Farrell**: Investigation, Writing - Review & Editing. **Nesrine Harroum**: Investigation, Writing - Review & Editing. **Younes Bakhta**: Investigation, Data curation. **Liora Cohen**: Investigation, Data curation. **Benjamin Pageaux**: Conceptualization, Methodology, Writing - Review & Editing, Visualization, Supervision. **Jason L. Neva**: Conceptualization, Methodology, Writing - Review & Editing, Visualization, Supervision.

## Data availability

The datasets generated and analyzed during the current study are available from the corresponding author upon reasonable request, and approval of the ethics committee. Individual data are presented in the figures.

## Supplementary Material

**Table S1:**
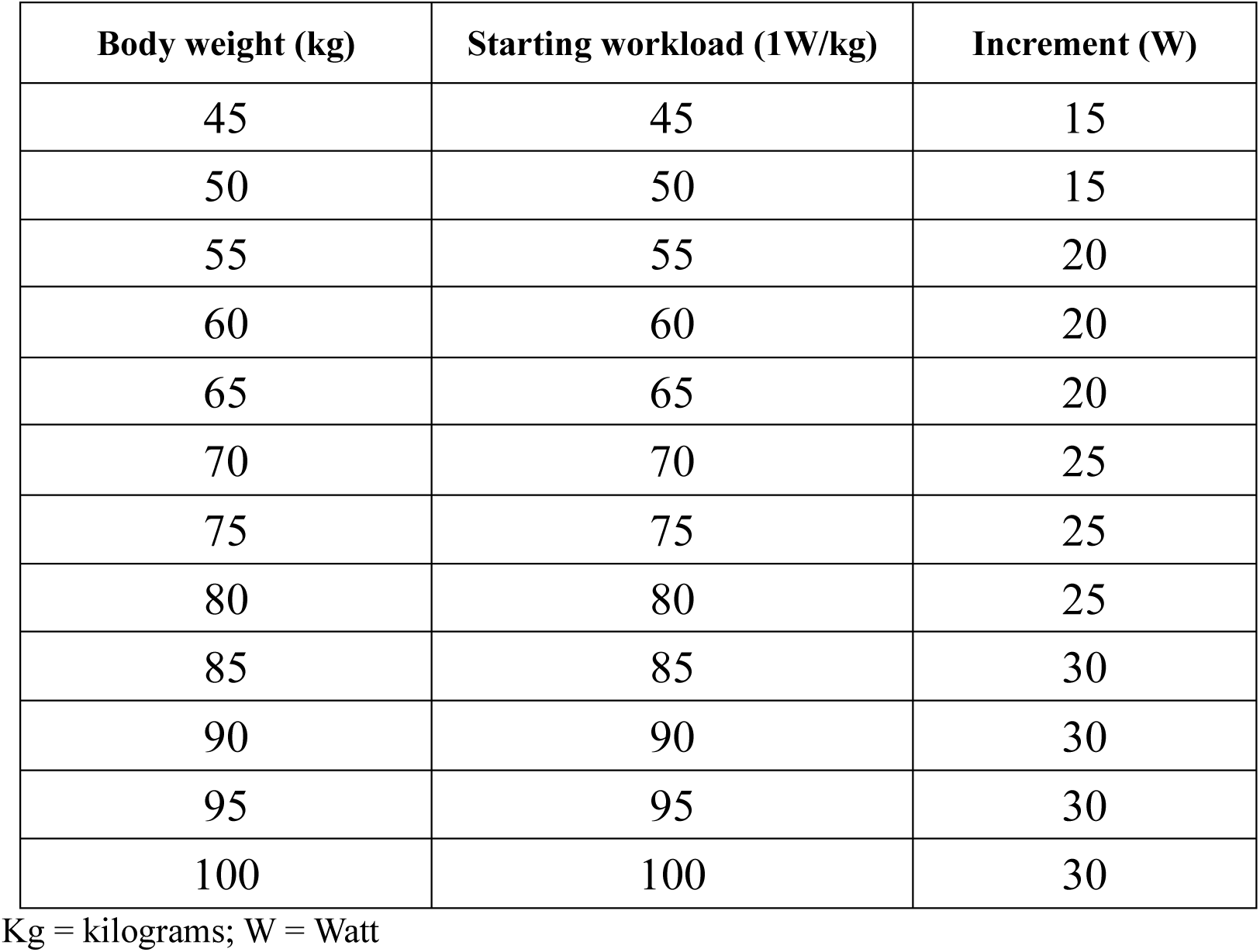
Starting workloads and increments for the incremental exercise test.

**Table S2:** Resting motor threshold across experimental interventions.

| TMS Current (% MSO) |  |  |  | 880 |
| --- | --- | --- | --- | --- |
| Condition | LM | PA | AP | 881 |
| Eccentric | 54.63 ± 7.94 | 54.00 ± 7.31 | 66.40 ± 9.50 | 882 |
| Concentric | 56.43 ± 9.10 | 54.40 ± 7.47 | 66.53 ± 8.97 | 883 |
| Rest | 56.47 ± 7.61 | 53.67 ± 7.57 | 66.27 ± 8.21 | 884 |

Results are presented as mean ± SD. Abbreviations: TMS = transcranial magnetic stimulation; % MSO = percentage of maximum stimulator output; PA = posterior-to-anterior; AP = anterior-to-posterior; LM = lateral-to-medial.

**Table S3:** Test stimulus across experimental interventions and time-points.

|  |  | % MSO |  |  | MEP Amplitude |  |  |
| --- | --- | --- | --- | --- | --- | --- | --- |
| TMS current | Condition | T<sub>0</sub> | T<sub>1</sub> | T<sub>2</sub> | T<sub>0</sub> | T<sub>1</sub> | T<sub>2</sub> |
| PA | Eccentric | 71.47 ± 14.21 | 72.16 ± 14.17 | 70.13 ± 12.71 | 1.06 ± 0.11 | 1.10 ± 0.11 | 1.10 ± 0.11 |
|  | Concentric | 73.73 ± 15.23 | 72.87 ± 13.81 | 72.10 ± 14.87 | 1.07 ± 0.11 | 1.08 ± 0.12 | 1.08 ± 0.11 |
|  | Rest | 70.60 ± 11.33 | 72.10 ± 12.54 | 70.43 ± 13.38 | 1.10 ± 0.09 | 1.13 ± 0.07 | 1.11 ± 0.09 |
| AP | Eccentric | 83.77 ± 12.6 | 85.43 ± 12.13 | 84.76 ± 11.05 | 1.06 ± 0.17 | 1.04 ± 0.18 | 1.09 ± 0.17 |
|  | Concentric | 84.73 ± 12.07 | 86.23 ± 12.26 | 85.46 ± 12.36 | 1.04 ± 0.19 | 1.07 ± 0.19 | 1.06 ± 0.19 |
|  | Rest | 85.83 ± 11.41 | 84.97 ± 13.81 | 83.66 ± 13.41 | 1.06 ± 0.15 | 1.07 ± 0.15 | 1.08 ± 0.15 |

Results are presented as mean ± SD. Abbreviations: TMS = transcranial magnetic stimulation; % MSO = percentage of maximum stimulator output; PA = posterior-to-anterior; AP = anterior-to-posterior; T<sub>0</sub> = before the experimental intervention, T<sub>1</sub> = immediately after the experimental intervention, T<sub>2</sub> = 20 min after the experimental intervention.

**Figure S1.**
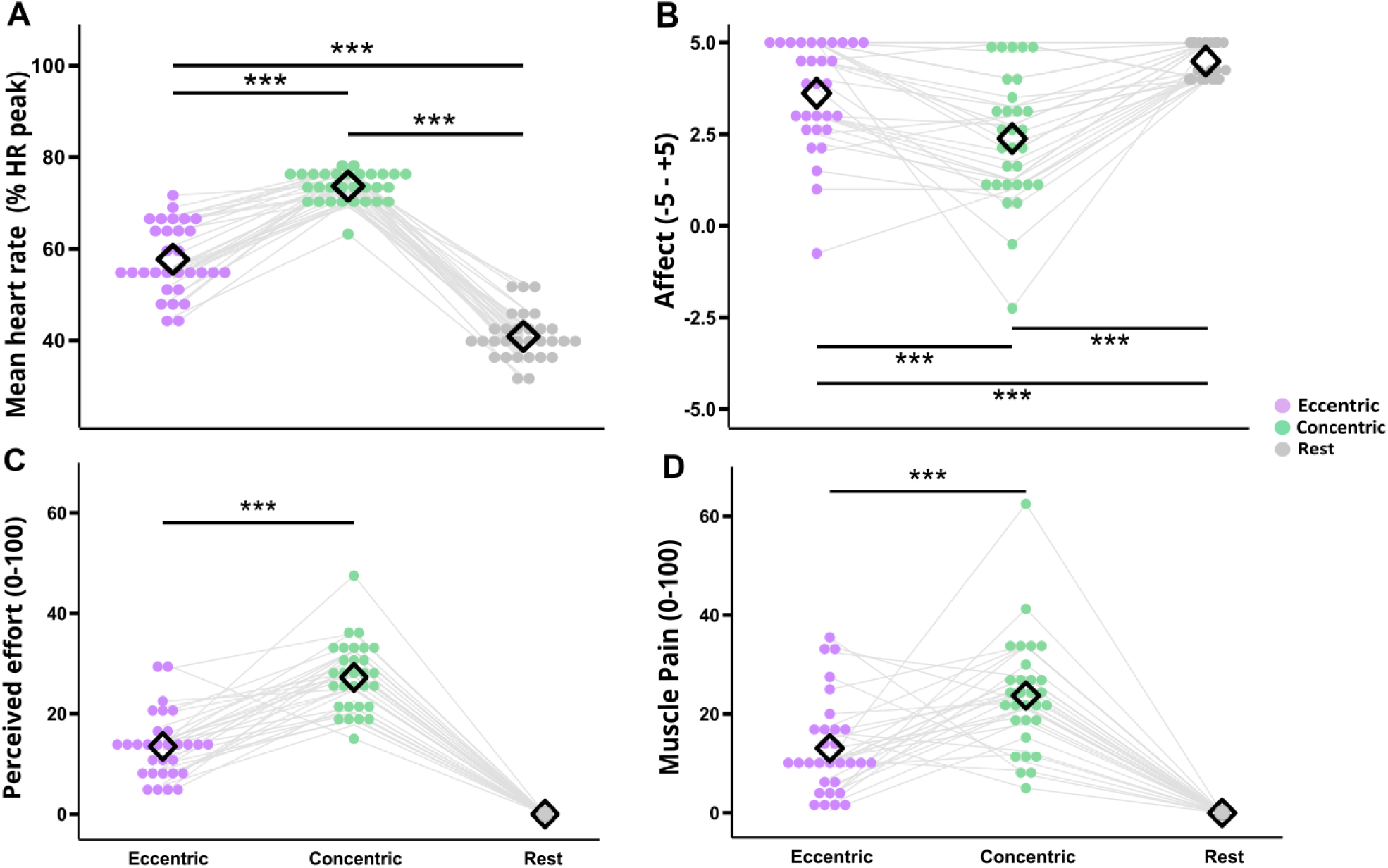
Parameters obtained during the different experimental interventions. All parameters are averaged over the 20-minute intervention (eccentric cycling, concentric cycling and rest). Panel A: Mean heart rate (%HRpeak). Panel B: affect (feeling scale from -5 to +5). Panel C: perceived effort (CR100 scale). Panel D: muscle pain (CR100 scale). Mean heart rate during the 20-min intervention was expressed relative to HRpeak obtained during the concentric incremental test. Dots represent individual data points, while the black diamond represents the mean for each condition. Purple dots refer to the eccentric cycling condition, green dots to the concentric cycling condition, and grey dots to the rest condition. Grey lines connect the data points across the three conditions for the same participants. ***: *p*-value < .001. For perceived effort and muscle pain, the rest condition was excluded from the analysis as all participants reported a score of 0, allowing for a direct comparison between eccentric and concentric cycling. HRpeak = peak heart rate.

**Figure S2.**
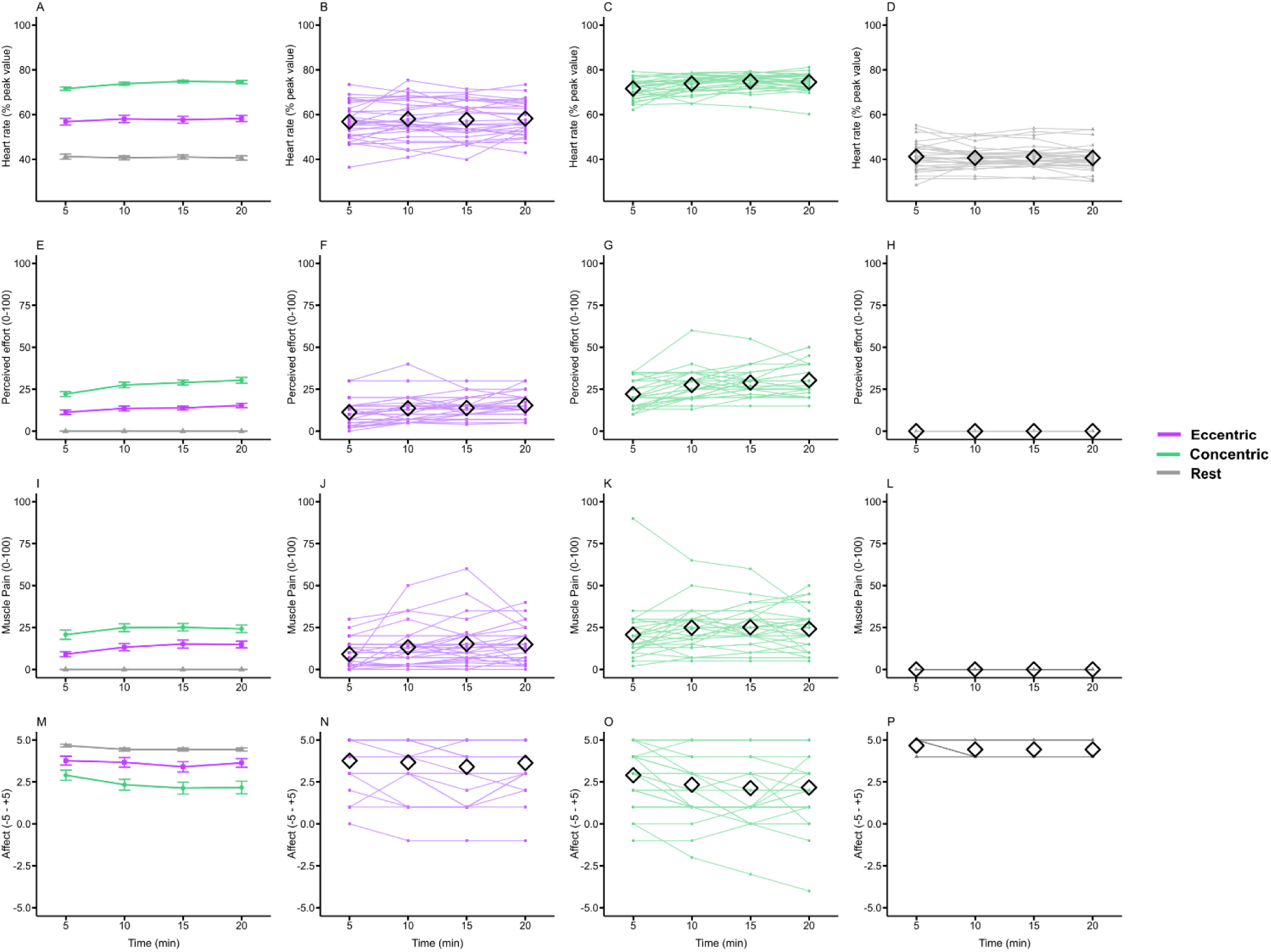
Parameters obtained across the different experimental interventions. Green represents concentric condition, purple represents eccentric condition, and grey represents rest condition. Time points of 5, 10, 15, and 20 minutes indicate when parameters were recorded during the intervention. The black diamond shape indicates the mean at each time point. Heart rate (% peak value): Panel A shows the means for each condition, while Panels B, C, and D display individual data points for the concentric, eccentric, and rest conditions, respectively. Perceived effort: Panel E shows the means for each condition, while Panels F, G, and H display individual data points for the concentric, eccentric, and rest interventions, respectively. Muscle pain: Panel I shows the means for each condition, while Panels J, K, and L display individual data points for the concentric, eccentric, and rest conditions, respectively. Affect: Panel M shows the means for each group, while Panels N, O, and P display individual data points for the concentric, eccentric, and rest conditions, respectively.

**Figure S3.**
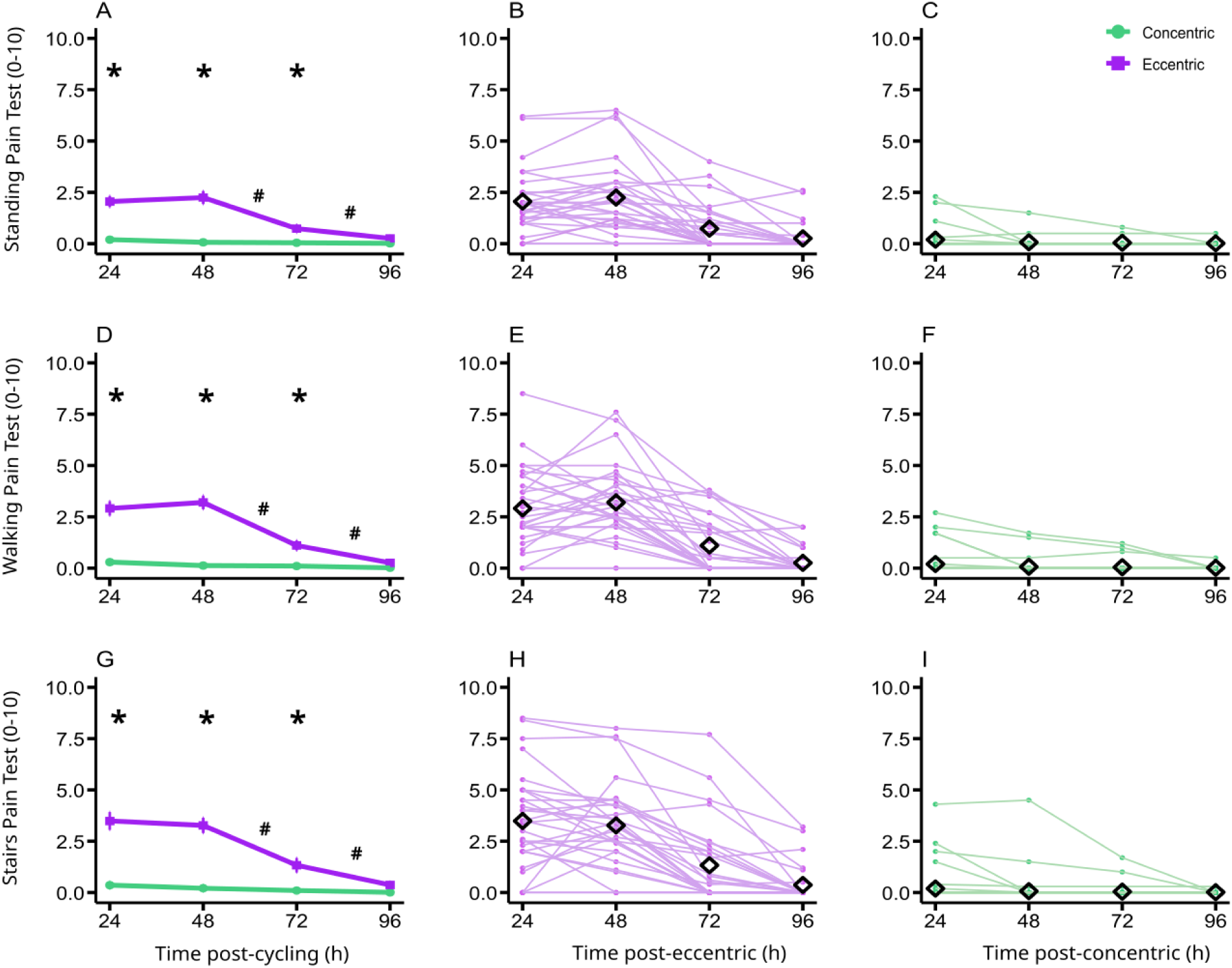
Muscle pain assessments after cycling exercise. Purple represents eccentric condition, green represents concentric condition. Time points of 24, 48, 72, and 96 hours indicate when pain was assessed following the exercise intervention. The black diamond shape indicates the mean at each time point. Bars represent the standard error of the mean. *Standing pain test:* Participants were asked to rate their muscle pain when standing without any movement; Panel A shows the means for each condition, while Panels B and C display individual data points for the eccentric and concentric conditions, respectively. *Walking pain test:* Participants were asked to rate their muscle pain when walking; Panel D shows the means for each condition, while Panels E and F display individual data points for the eccentric and concentric conditions, respectively. *Stairs pain test:* Participants were asked to rate their muscle pain when taking the stairs up and down; Panel G shows the means for each condition, while Panels H and I display individual data points for the eccentric and concentric conditions, respectively. * indicates significant difference between both cycling exercise conditions at the specific time point (*p-value* < .05). # indicates a significant difference between the two adjacent time points (*p- value* < .05).

**Figure S4.**
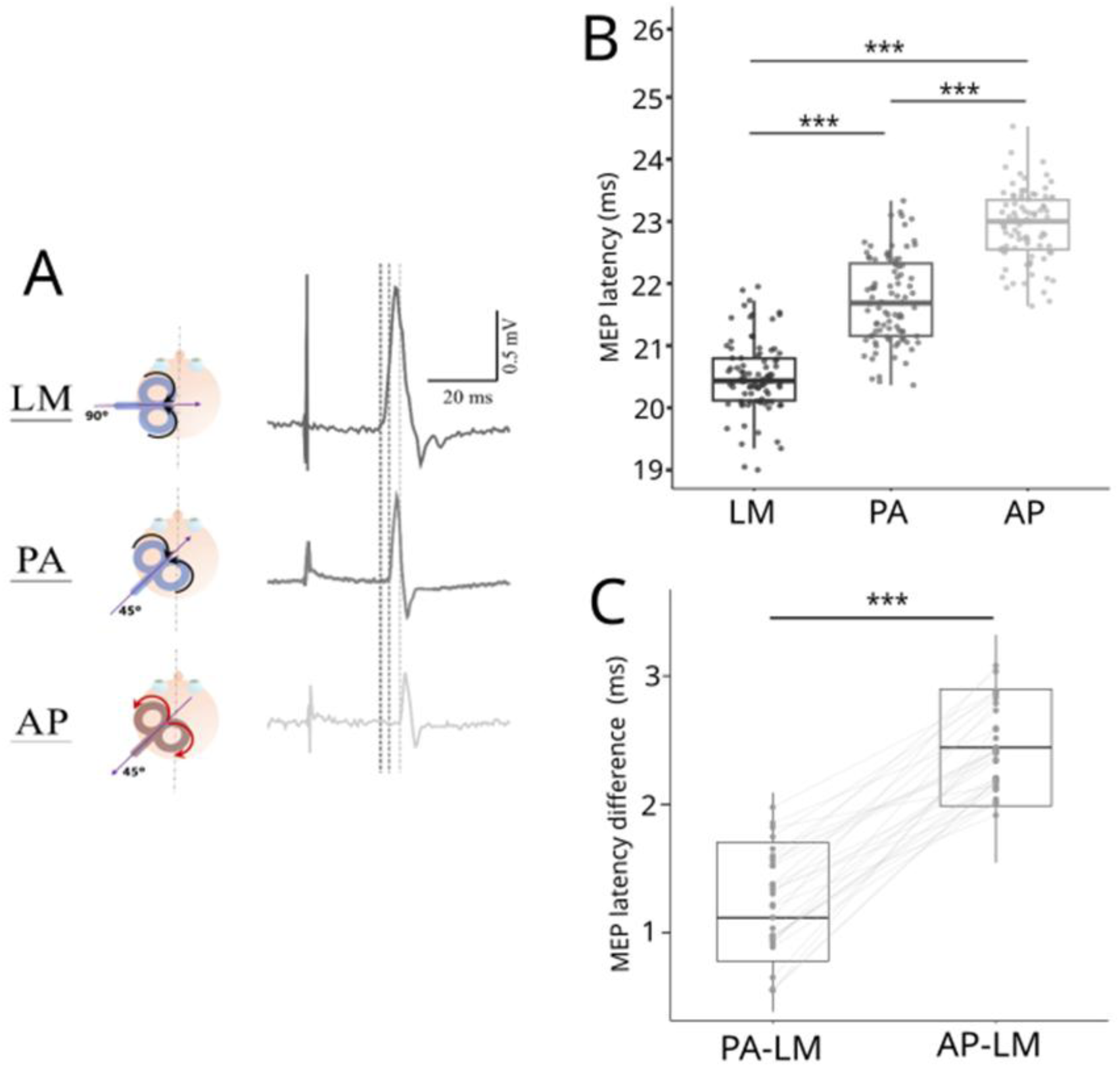
Transcranial magnetic stimulation (TMS) current directions and motor evoked potential (MEP) onset latency results. Panel A displays TMS current directions. LM TMS is shown in black, PA TMS in dark grey, and AP TMS in light grey. The figure illustrates the TMS coil current directions, depicted by purple arrows, and their orientations over the left M1 (dominant) abductor pollicis brevis (APB) muscle representation. A standard 8-figure TMS coil was used for both LM and PA stimulations, and a reversed current coil for AP stimulations. The coil was oriented at 90° for LM TMS and at 45° for both PA and AP TMS, relative to the longitudinal fissure. Electromyographic (EMG) traces from a representative participant, recorded from the right (dominant) APB are shown. Vertical dotted lines represent MEP onset latency elicited by the different TMS current directions (LM, PA, AP). Panel B displays boxplots of MEP onset latency for each TMS current (LM, PA, AP), with dots representing individual data. Panel C displays MEP onset latency differences (LM-PA, LM-AP) using boxplots, with connected individual data. AP = anterior-to-posterior; LM = lateral-to-medial; ms: milliseconds; PA = posterior-to-anterior; TMS = transcranial magnetic stimulation; *** *p* < .001.

**Figure S5.**
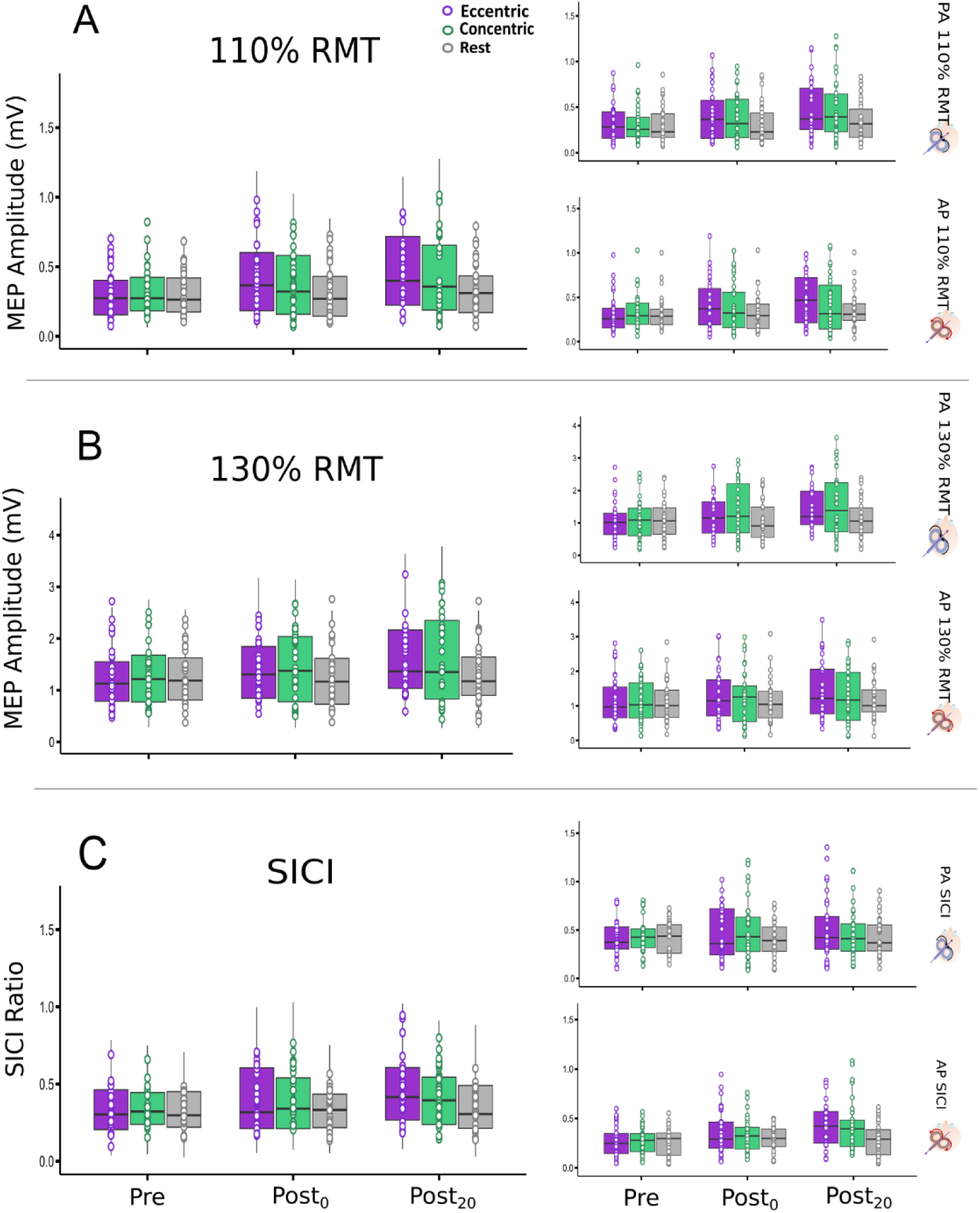
Individual data for neurophysiological measures. Panels A and B display boxplots of peak-to-peak MEP amplitudes for 110% RMT and 130% RMT respectively, and Panel C represents boxplots for average SICI ratios with TMS currents collapsed et each time point. Subplots representing PA and AP currents for each measure are displayed on the right. Purple color refers to eccentric condition, green color refers to concentric condition and grey color refers to rest condition. TMS = transcranial magnetic stimulation. AP = anterior-to-posterior; PA = posterior-to-anterior; MEP: motor-evoked potential; RMT = resting motor threshold; SICI = short-interval intracortical inhibition; mV: millivolt; Pre: before the experimental intervention; Post_0_: immediately after the experimental intervention; Post_20_: 20 minutes after the experimental intervention.

**Figure S6.**
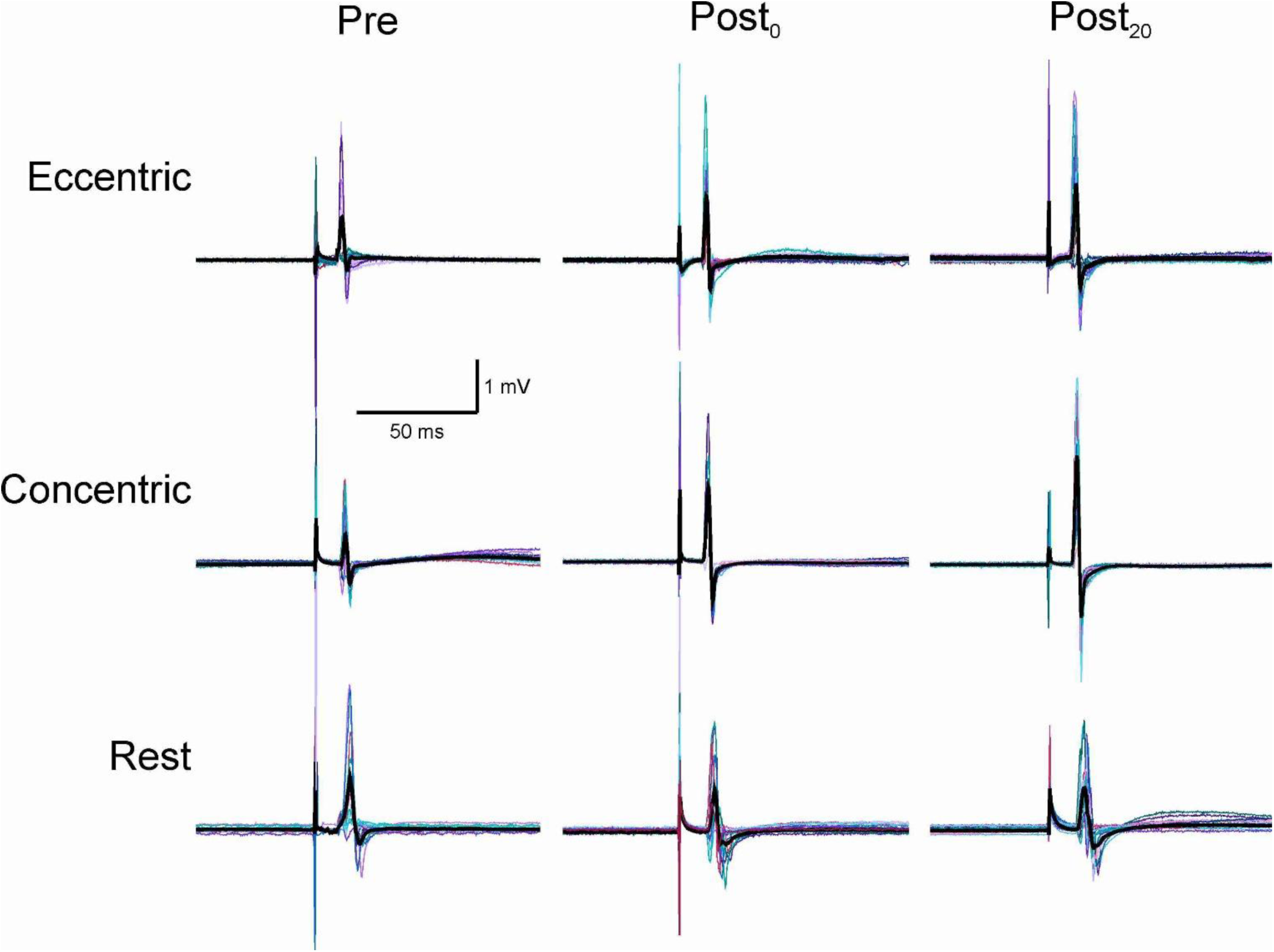
Individual MEP traces for 130% RMT. Representative motor-evoked potential (MEP) traces from a single participant under each experimental condition collected at three time points: Pre (before cycling exercise/rest), Post_0_ (immediately after cycling exercise/rest) and Post_20_ (20 min after cycling exercise/rest). Fifteen MEP trials were performed at each time point, with the black trace representing the average waveform of the 15 trials.

